# Cross-species modeling of plant genomes at single nucleotide resolution using a pre-trained DNA language model

**DOI:** 10.1101/2024.06.04.596709

**Authors:** Jingjing Zhai, Aaron Gokaslan, Yair Schiff, Ana Berthel, Zong-Yan Liu, Wei-Yun Lai, Zachary R. Miller, Armin Scheben, Michelle C. Stitzer, M. Cinta Romay, Edward S. Buckler, Volodymyr Kuleshov

## Abstract

Interpreting function and fitness effects in diverse plant genomes requires transferable models. Language models (LMs) pre-trained on large-scale biological sequences can learn evolutionary conservation and offer cross-species prediction better than supervised models through fine-tuning limited labeled data. We introduce PlantCaduceus, a plant DNA LM based on the Caduceus and Mamba architectures, pre-trained on a curated dataset of 16 Angiosperm genomes. Fine-tuning PlantCaduceus on limited labeled Arabidopsis data for four tasks, including predicting translation initiation/termination sites and splice donor and acceptor sites, demonstrated high transferability to 160 million year diverged maize, outperforming the best existing DNA LM by 1.45 to 7.23-fold. PlantCaduceus is competitive to state-of-the-art protein LMs in terms of deleterious mutation identification, and is threefold better than PhyloP. Additionally, PlantCaduceus successfully identifies well-known causal variants in both Arabidopsis and maize. Overall, PlantCaduceus is a versatile DNA LM that can accelerate plant genomics and crop breeding applications.

## Main

Over 1,000 plant genomes have been published during the past 20 years, and this number will continue to increase significantly in the coming decades ^1–3^. Understanding the functional elements and fitness effects of these genomes at both transcriptional and translational levels is crucial for advancing plant genomics and crop breeding. Unlike biomedical applications that primarily focus on a few key species, plant genomics must account for the vast diversity of hundreds of crop species, each with unique variations in size, composition, and complexity ^4^. Extensive genomic resources have been generated for model plants, such as Arabidopsis ^5^, rice ^6^ and maize ^6^, significantly advancing plant genomics research. However, generating analogous genomic resources experimentally for all plant genomes is time-consuming, costly, and impractical. This highlights the need for developing cross-species models capable of capturing evolutionary conservation across diverse plant species.

Supervised deep learning (DL) sequence models are successful in understanding DNA sequence functions such as transcription initiation ^7^, alternative splicing ^8^ and gene expression ^9^. However, supervised DL models typically require large-scale labeled data, such as ENCODE-scale datasets ^10,11^, to achieve robust performance. Such extensive labeled data is often scarce in plant genomics. Moreover, training supervised models on model species, such as Arabidopsis, presents challenges when transferring to other plant species. However, the success of self-supervised language models (LMs) offers a promising alternative. In this paradigm, a foundation model is pre-trained on vast amounts of unlabeled biological sequences to learn evolutionary conservation. Pre-trained models are then fine-tuned on limited labeled data, enabling better performance on downstream tasks and enhancing generalizability across species relative to existing methods. For example, protein LMs, pre-trained on diverse protein sequences spanning the evolutionary tree, have shown successful applications in predicting atomic-level protein structure ^12^ and disease-causing variants ^13^ as well as in engineering protein design ^14^. These models provide valuable tools for understanding protein function and facilitating innovative solutions in biotechnology and medicine ^15^.

Unlike protein LMs that are limited to coding regions, DNA LMs enable a comprehensive understanding of the entire genome, offering deeper insights into gene regulation and evolution. Protein LMs have shown success in identifying pathogenic missense mutations in human genetics^13,16^, but increasing evidence shows that mutations in noncoding regions, including both intergenic and intronic regions, contribute significantly to both agronomic traits ^17^ and human diseases ^18,19^. Additionally, training multi-species DNA LMs can capture evolutionary conservation at the DNA level, enhancing our understanding of genetic variation across different species.

However, DNA LMs face significant challenges compared to protein LMs. Firstly, eukaryotes, especially plants ^20^, contain varied percentages of repetitive sequences, complicating the pre-training task. Given that LMs are pre-trained to either predict the next token or tokens are masked arbitrarily in a sequence, repetitive sequences that are easier to predict but do not necessarily improve downstream applications can reduce overall model quality ^21^. Additionally, noncoding regions are less conserved than coding regions, leading to potential biases if entire genomes are included in pre-training. Lastly, unlike protein sequences, modeling double-stranded DNA requires consideration of reverse complementary base pairing ^22^ and a bi-directional model that accounts for both upstream and downstream sequences.

To tackle these challenges, we introduce PlantCaduceus, a DNA language model pre-trained on a curated dataset consisting of 16 angiosperm genomes (**Fig. 1A-1B**). PlantCaduceus employs single-nucleotide tokenization, enabling precise modeling at the base-pair-resolution across diverse plant genomes. By down-sampling noncoding regions and down-weighting repetitive sequences, we generated an unbiased genomic dataset for pre-training. In contrast, other publicly available DNA LMs, such as AgroNT ^23^ and Nucleotide Transformer ^24^, use entire genomes for pre-training, potentially introducing biases toward certain genomes and repetitive sequences. Additionally, both models use non-overlapping kmer tokenizers that disrupt the genome into arbitrary segments. Unlike the unidirectional HyenaDNA ^25^ or Evo ^26^, PlantCaduceus offers bi-directional context, providing a more comprehensive understanding of DNA interactions. Furthermore, to handle double-stranded DNA, we used the Caduceus architecture ^27^, which builds on the Mamba architecture ^28^ and supports reverse complement equivariance, unlike GPN ^21^, which uses convolutional neural network and manually augments reverse complement sequences. By evaluating the pre-trained PlantCaduceus model on five cross-species tasks, including translation initiation/termination sites, splice donor and acceptor sites, and evolutionary conservation prediction. We found that our model demonstrated the best performance compared to baseline models for all five tasks. Notably, downstream classifiers fine-tuned on PlantCaduceus with limited labeled data in Arabidopsis maintained the best performance on other crop species such as maize, improving the PRAUC from 1.45-fold to 7.23-fold as compared to the best existing DNA LM, indicating that PlantCaduceus effectively captures broad evolutionary conservation. Additionally, deleterious mutations identified with the zero-shot strategy of PlantCaduceus showed a three-fold enrichment of rare alleles when compared to the most commonly used evolutionary-based methods such as phyloP and phastCons ^29^. For missense mutations, PlantCaduceus matches the performance of state-of-the-art protein LMs, suggesting that PlantCaduceus can be effectively used for genome-wide deleterious mutation identification. Furthermore, PlantCaduceus successfully identifies well-known causal variants in both Arabidopsis and maize. These results indicate that PlantCaduceus can serve as a foundational model to accelerate plant genomics and crop breeding applications.

**Fig 1.**
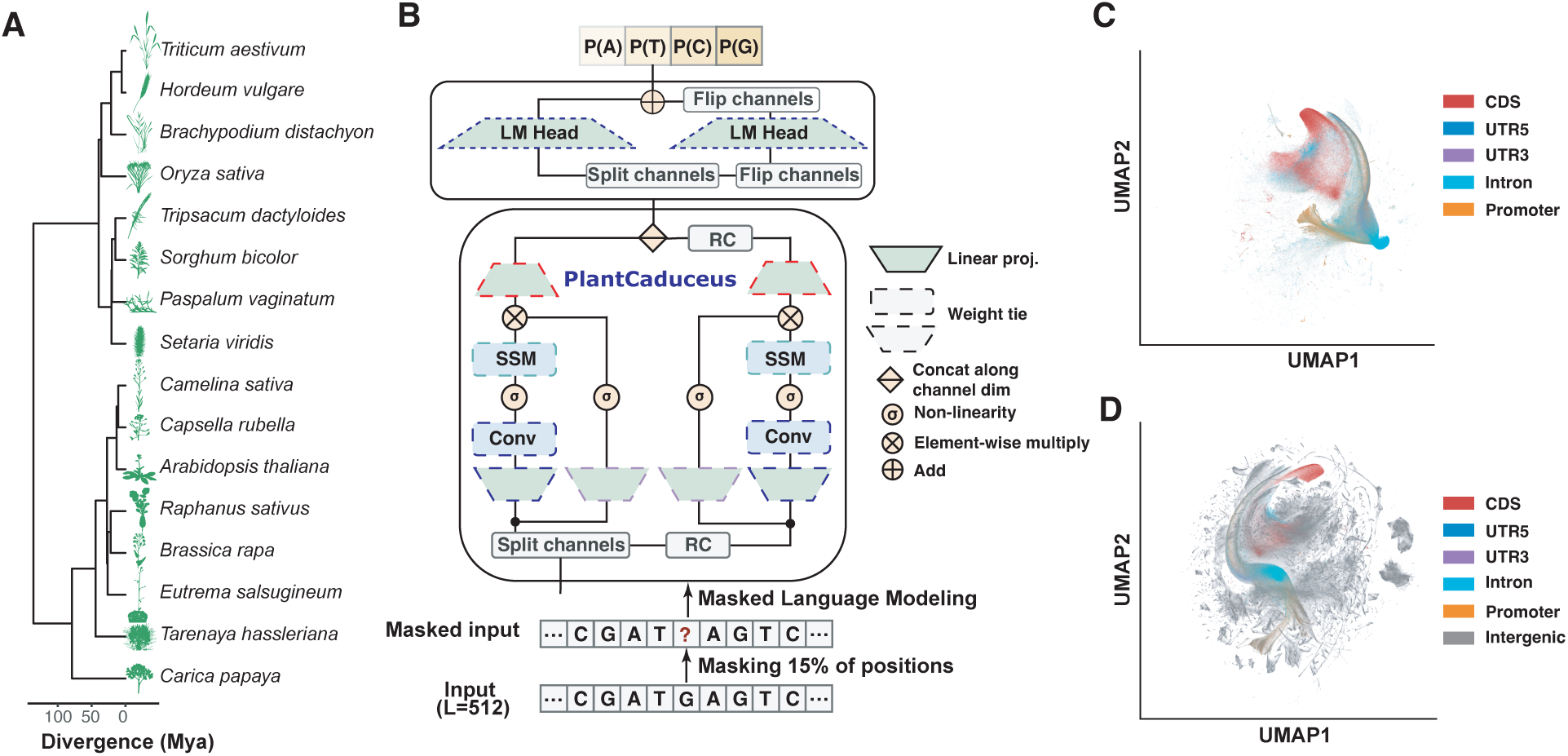
Overview of PlantCaduceus. **(A)** Phylogenetic tree of 16 Angiosperm species used for pre-training the PlantCaduceus model. **(B)** The input for PlantCaduceus consists of 512-bp DNA sequences with 15% of positions randomly masked. The pre-training objective is cross-entropy loss on the masked positions. The sequences are processed through the bi-directional Caduceus architecture, which is based on the Mamba sequence operator—a recently proposed structured state space model. Caduceus also contains a reverse complement equivariance inductive bias. **(C)** UMAP visualization of embeddings from PlantCaduceus (32 layers) averaged over non-overlapping 100-bp windows along the sorghum genome without intergenic regions. **(D)** The same UMAP visualization as in **(C)** but with intergenic regions.

## Results

### PlantCaduceus: a pre-trained DNA language model with 16 Angiosperm genomes

Caduceus ^27^ is a DNA LM architecture that builds upon the recently introduced Mamba ^28^ architecture, a selective state space sequence model that has demonstrated competitive performance to transformers ^30^ in various natural language processing tasks, with more efficient scaling for longer range sequences. Unlike Mamba, Caduceus is specifically designed for DNA sequences, taking into account the bi-directional nature of DNA and introducing reverse complement (RC) equivariance. Here, we trained PlantCaduceus using the Caduceus architecture on 16 Angiosperm genomes (**Fig. 1A-1B; Supplemental Table 1**), spanning 160 million years of evolutionary history (**METHODS**). PlantCaduceus takes 512 base pair (bp) windows of input sequences, tokenizing them into single nucleotides, and is pre-trained using a masked language modeling objective (**Fig. 1B; METHODS**). To address the substantial variation in genome sizes and the high proportion of repetitive sequences in these genomes, we emphasized non-repetitive sequences by down-weighting and down-sampling repetitive sequences during pre-training (**METHODS**). To scale Caduceus, we trained a series of PlantCaduceus models with parameter sizes ranging from 20 million to 225 million (**Table 1**). The training and validation losses for each model are detailed in **Supplemental Table 2**. After pre-training, we conducted a preliminary assessment to verify the model’s learning capabilities. Taking the sorghum genome as an example, we employed Uniform Manifold Approximation and Projection (UMAP) ^31^ to visualize the embeddings generated by the four pre-trained PlantCaduceus models. By segmenting the genome into 512 bp windows, we observed distinct clustering in the UMAP visualization, corresponding to different genomic regions (**Fig. 1C**). Due to the high proportion of repetitive intergenic sequences in the sorghum genome, the embedding spaces appeared dispersed in the UMAP visualization (**Fig. 1D**; **Supplemental Fig. 1**). Even without any supervision, PlantCaduceus was able to differentiate between coding and noncoding regions with high clarity.

**Table 1.**
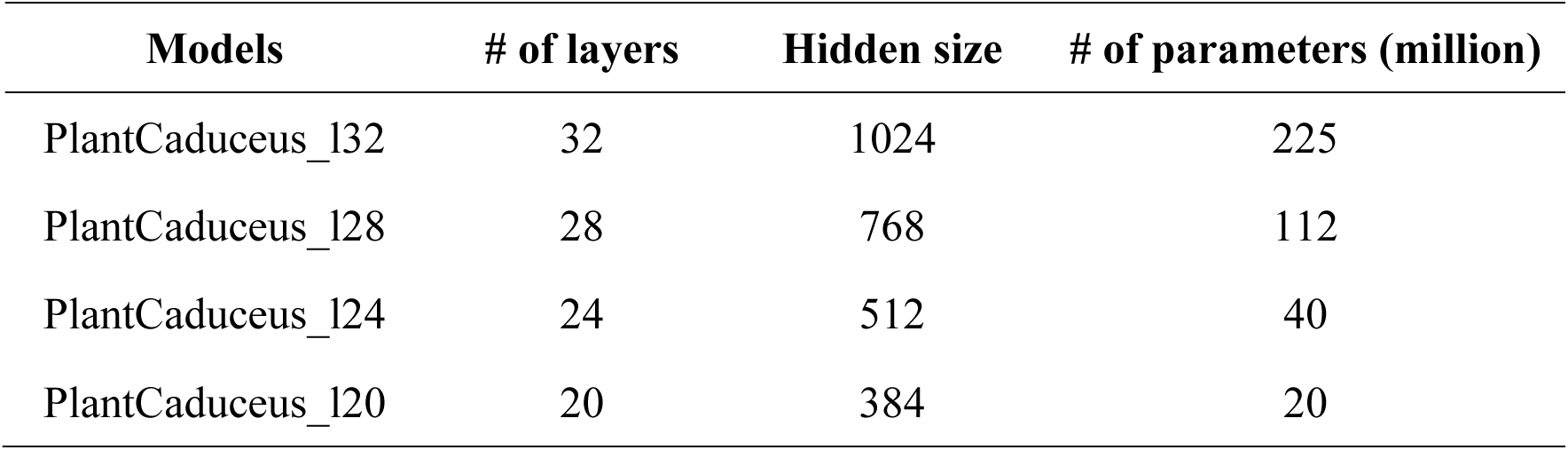
PlantCaduceus model parameters.

### Improving the accuracy and cross-species transferability of modeling transcription and translation through fine-tuning PlantCaduceus

Transcription and translation are two key processes in the central dogma of molecular biology, and the precise identification of junction sites during these processes is essential for comprehensive gene annotation. To assess PlantCaduceus’s performance in modeling these processes, we designed four gene annotation tasks: predicting the translation initiation site (TIS), translation termination site (TTS), and splice donor and acceptor sites (**METHODS**). We employed a feature-based approach to fine-tune PlantCaduceus by keeping the pre-trained model weights frozen while training XGBoost models using embeddings extracted from the last hidden state of PlantCaduceus (**Fig. 2A**). Compared to full fine-tuning, this approach allows us to leverage the rich representations learned by PlantCaduceus while minimizing the usage of computational resources. Previous LMs focus on evaluation within the same species ^24,25,32–34^. However, given that the DNA LM model is pre-trained on multiple species, we wanted to investigate whether a model fine-tuned with limited labeled data in Arabidopsis could be used for prediction in other species. Therefore, we trained and validated XGBoost models in Arabidopsis and tested their performance on both species included (*Oryza sativa* and *Sorghum bicolor*) and not included (*Gossypium hirsutum*, *Glycine max* and *Zea mays*) in the pre-training (**Fig. 2B; Supplemental Table 3**). We benchmarked the performance of PlantCaduceus against three DNA LMs: GPN ^21^, AgroNT ^23^, and Nucleotide Transformer ^24^, as well as a supervised hybrid model comprising a convolutional neural network (CNN) and a long short-term memory (LSTM) network ^35^, hereafter referred to as CNN+LSTM. For DNA LMs, we used the same feature-based approach as PlantCaduceus (**Fig. 2A**) to train XGBoost models using embeddings extracted from the last hidden state of each DNA LM (**Fig. 2C**). The CNN+LSTM model was trained from scratch in a supervised manner.

**Fig 2.**
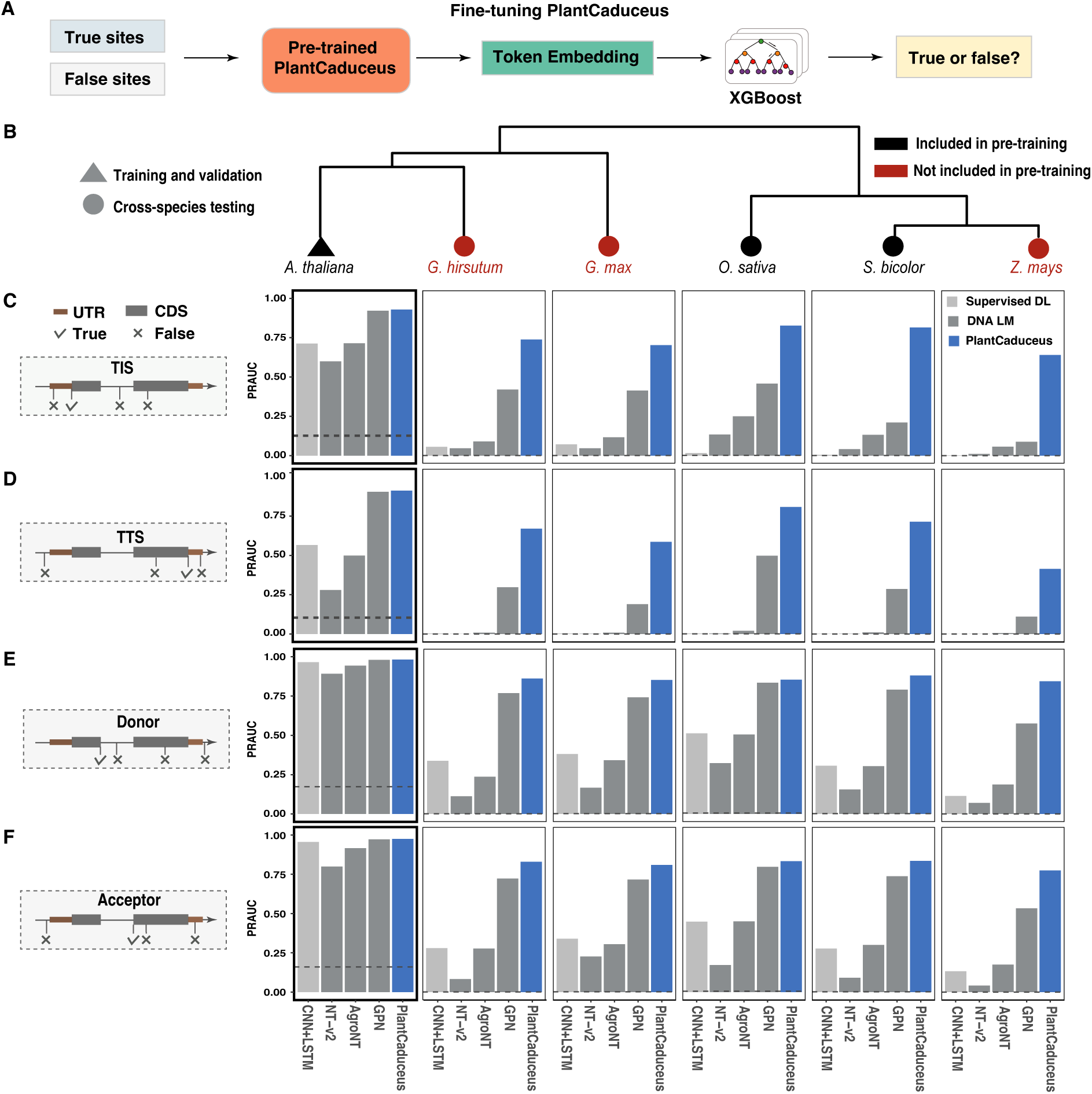
Modeling translation and transcription through fine-tuning PlantCaduceus. **(A)** Fine-tuning strategy for PlantCaduceus: The weights of the pre-trained PlantCaduceus model are kept frozen during pre-training. The last hidden state of PlantCaduceus is then used as features for the XGBoost model. **(B)** Phylogenetic tree of species used for training, validation, and testing during the fine-tuning of PlantCaduceus. **(C-F)** Bar plots displaying the PRAUC scores for six species across four tasks: TIS **(C)**, TTS **(D)**, splice donor **(E)**, and splice acceptor **(F)**. The gene structures on the left illustrate how positive and negative samples are obtained for each classification task. Blue bars represent the PlantCaduceus model with 32 layers. Gray bars denote three DNA language models: NT-v2, AgroNT, and GPN. Light gray bars represent a traditional supervised model, a hybrid of CNN and LSTM. The gray dashed line in each panel indicates the baseline for each dataset, corresponding to the negative sample ratio.

First, focusing on within species evaluation on Arabidopsis hold-out test set, PlantCaduceus (32 layers) showed consistently superior performance across the four tasks of predicting TIS (**Fig. 2C**), TTS (**Fig. 2D**), splice donor site (**Fig. 2E**), and splice acceptor site (**Fig. 2F**). Other DNA LMs like GPN and AgroNT also performed well, particularly in predicting splice donor and acceptor sites. Additionally, for splice donor and acceptor site prediction, even the supervised CNN+LSTM model achieved near perfect PRAUC values, indicating that within-species prediction is a relatively straightforward task.

We then assessed the cross-species generalization ability of these models by testing them on *O. sativa* and *S. bicolor*, which were included in pre-training, as well as *G. hirsutum*, *G. max*, and *Z. mays*, which were not (**Fig. 2B**; **Fig. 1A**). When tested across these five species, all models except PlantCaduceus exhibited a significant drop in average PRAUC, decreasing from 0.789 in *A. thaliana* to 0.237 in these species (**Fig. 2C-2F**). For instance, transferring the supervised CNN+LSTM model to *Z.mays*—which diverged 160 million years ago—resulted in a PRAUC drop from 0.713 to nearly zero for the TIS task. This significant drop was expected, as the supervised model had never seen sequences from these species, making cross-species generalization challenging. Although GPN maintained decent cross-species predictions, it still showed significant performance drops, with the average PRAUC decreasing from 0.944 in *A. thaliana* to 0.509 in other species (**Fig. 2C-2F; Supplemental Table 4**). As expected, the non-plant DNA NT-v2 model performed poorly on these tasks due to the significant divergence between plant and animal genomes. Even though AgroNT was pre-trained on 48 plant genomes, its performance fell short of expectations in cross-species evaluations. In contrast, PlantCaduceus consistently maintained high PRAUC values across all species, with an average PRAUC of 0.764, regardless of whether the species were included in pre-training, demonstrating its superior generalization ability across diverse plant species (**Fig. 2C-2F; Supplemental Table 4**).

GPN, as the second-best DNA LM, was not pre-trained on any of the five testing species. To ensure a fairer comparison with GPN and to understand why PlantCaduceus achieved superior performance on these cross-species tasks, we conducted an ablation test by re-training a custom GPN model (**METHODS**) using the same datasets as PlantCaduceus and scaling it to 130 million parameters, on the same order of magnitude as PlantCaduceus. We observed that including more genomes in the pre-training and scaling model size significantly improved GPN’s cross-species predictability (**Supplemental Fig. 2**; **Supplemental Table 4**), especially for TIS and TTS tasks. This indicates that when more genomes are included during pre-training, the embeddings learned by DNA LMs are more general across species. However, PlantCaduceus still exhibited the best performance, indicating that its architecture is superior to that of GPN. Moreover, even with a parameter size of 20 million—6.5 times smaller than the custom 130 million GPN and 3.25x times smaller than the original GPN—PlantCaduceus still outperformed all models in predicting TIS, TTS, splice donor, and splice acceptor sites. These results demonstrate that PlantCaduceus not only captures broader evolutionary conservation features but also is more parameter-efficient than other DNA LMs.

### Cross-species evolutionary constraint prediction through fine-tuning PlantCaduceus

Genome-wide association studies (GWAS) have identified thousands of variants associated with complex traits ^36^. However, identifying causal variants is complicated by linkage disequilibrium (LD), as significant SNPs identified by GWAS are usually in LD with causal variants ^37^. In contrast, evolutionary constraint, as evidenced by DNA conservation across species, can identify potential causal mutations by revealing their fitness effects ^38^. Given that PlantCaduceus is pre-trained on 16 Angiosperm genomes, we hypothesize that it can be fine-tuned to predict evolutionary constraint using DNA sequences alone. Maize and sorghum are both members of the Andropogoneae clade, descended from a common ancestor approximately 18 million years ago ^39^. To generate evolutionary constraints in the sorghum genome, we aligned 34 genomes from the Andropogoneae clade, with rice as an outgroup (**Supplemental Table 5**), to the *Sorghum bicolor* reference genome (**Supplemental Fig. 3**). We focused on the 277 million sites with nearly complete coverage and defined those sites with an identity threshold of 15 as conserved versus neutral with an identity threshold of 15 (**Fig. 3A**). We used sites chromosomes 1 to 9 to train an XGBoost model and evaluated it on sorghum chromosome 10. As mentioned above, we benchmarked this task against GPN, AgroNT, NT-v2, and the supervised CNN+LSTM model. On the validation set, PlantCaduceus achieved the best performance, with an AUC of 0.896 (**Fig. 3B**) and a PR-AUC of 0.876 (**Fig. 3C**). In comparison, the best AUC and PR-AUC for other DNA LMs were 0.778 and 0.790, respectively. As expected, the supervised CNN+LSTM model performed the worst, with an AUC of 0.638, as it had only seen sequences from sorghum (**Fig. 3B-3C**). This demonstrates that PlantCaduceus enables predicting evolutionary constraint without multiple sequence alignment.

**Fig 3.**
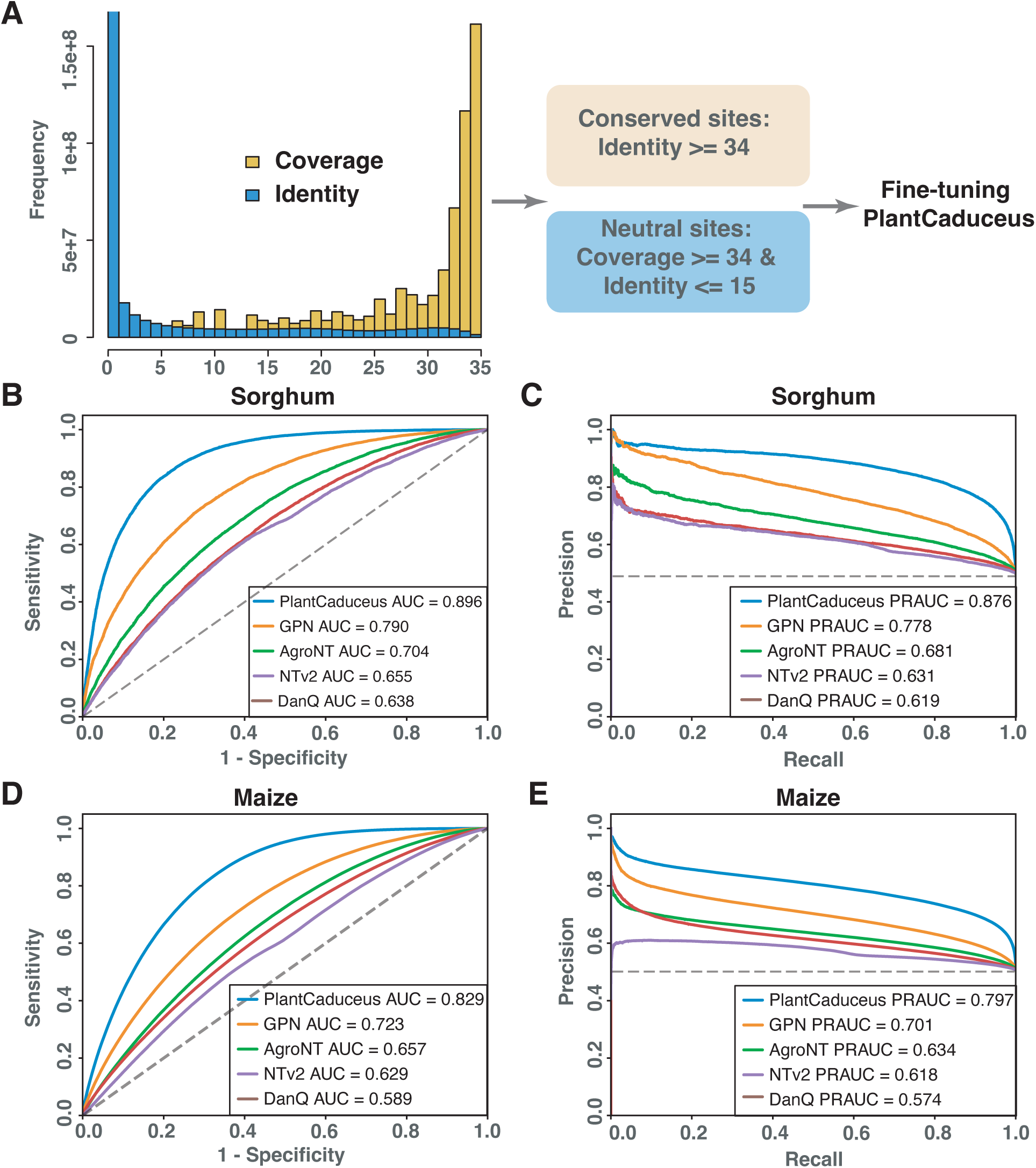
Evolutionary constraint prediction. **(A)** Illustration of the evolutionary conservation data curation. **(B)** Receiver operating characteristic (ROC) and **(C)** precision-recall (PR) curves of different models in sorghum. **(D)** ROC and **(E)** PR curves of transferring different models trained in sorghum to unseen maize data.

To further explore the cross-species predictive power of the model fine-tuned on sorghum evolutionary constraint data, we generated an analogous testing dataset for maize (**METHODS**). Remarkably, when our PlantCaduceus model, originally fine-tuned on sorghum, was applied to the maize dataset, it demonstrated strong cross-species prediction performance, achieving an AUC of 0.829 (**Fig. 4D**) and a PR-AUC of 0.797 (**Fig. 4E**). In contrast, all other models consistently showed poor performance on maize (**Fig. 4D-4E**). We also evaluated the performance of our custom GPN model which was trained on the same dataset as PlantCaduceus. While the custom GPN model showed improved performance with an AUC of 0.833 and a PR-AUC of 0.814, PlantCaduceus, with only 20 million parameters, outperformed both the original GPN and the custom GPN models (**Supplemental Fig.** 4). These results highlight the robustness and effectiveness of our DNA LM for cross-species predictions of evolutionary constraints using only sequence data as input. The transferability of our model across different species within the Andropogoneae clade suggests that it captures fundamental evolutionary patterns and can be readily adapted to predict evolutionary constraint in related species with limited additional training data.

**Fig 4.**
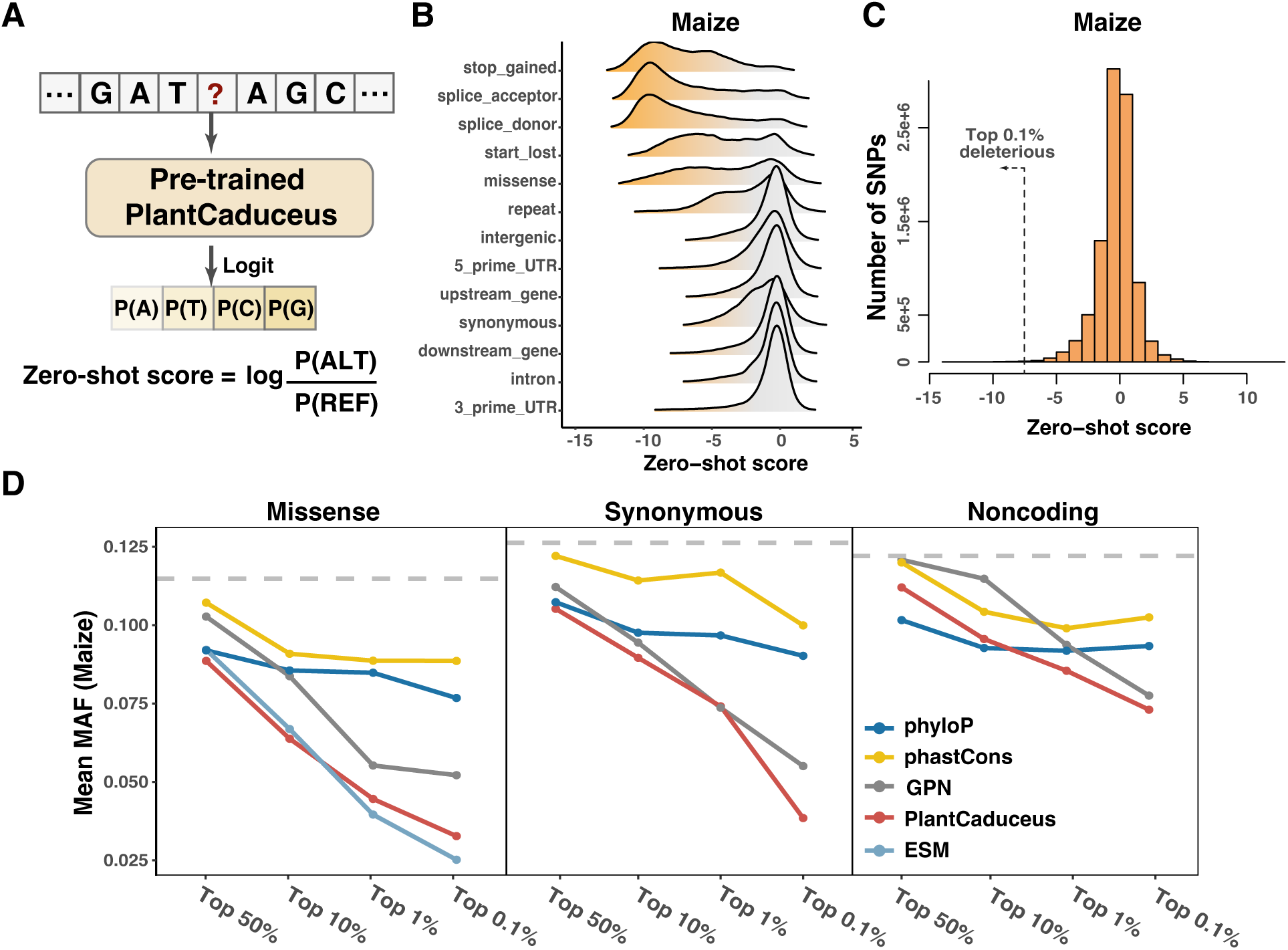
Deleterious mutations identification in maize. **(A)** The zero-shot strategy of PlantCaduceus for identifying deleterious mutations. **(B)** The zero-shot score distribution of different types of variants generated by in silico mutagenesis in maize chromosome 8. **(C)** The zero-shot score distribution of 9.4M SNPs in the maize Hapmap3 population. **(D)** The MAF of putative deleterious mutations prioritized by different models in maize.

### Zero-shot variant effect prediction identifies deleterious mutations in different species

The training objective of PlantCaduceus is to predict masked nucleotides based on sequence context; if a pre-trained multi-species DNA LM can accurately predict masked tokens, it suggests that similar sequence patterns, conserved across different species, were frequently observed during pre-training. We hypothesize that the predicted likelihood of the reference allele versus the alternate allele can identify deleterious mutations, as mutations in conserved regions across species are likely deleterious ^40–43^. To test this hypothesis, we employed the same zero-shot strategy as GPN ^21^ to estimate the effect of each mutation (**Fig. 4A**). Specifically, for each mutation, we calculated the log-likelihood difference between the reference and alternate alleles, where a more negative value indicates higher conservation. We generated 1.1 million sites from sorghum chromosome 8 (included in pre-training) and 1.3 million sites from maize chromosome 8 (not included) through in silico mutagenesis of SNPs. We then calculated zero-shot scores for these mutations to assess how PlantCaduceus performs on both seen and unseen genomes. As expected, mutations in highly conserved functional regions—such as stop-gained, splice acceptor, splice donor, and start-lost sites—exhibited the most negative zero-shot scores, underscoring their potential deleterious effects (**Fig. 4B; Supplemental Fig. 5A**). Missense mutations also showed notably negative zero-shot scores. In contrast, intergenic regions and introns displayed scores closer to zero, indicating lower evolutionary constraint and a reduced likelihood of deleterious effects (**Fig. 4B; Supplemental Fig. 5A**). However, we observed that a subset of mutations in repetitive regions still received very low zero-shot scores, suggesting that repetitive regions may be too easy for the model to predict the masked tokens. Overall, the zero-shot score of PlantCaduceus aligns with established concepts of deleteriousness ^44,45^.

Besides in silico mutagenesis, we also evaluated if zero-shot score can be used to identify deleterious mutations in natural populations. Deleterious mutations tend to have lower frequencies within a population due to selective constraints ^38^, we therefore used minor allele frequency (MAF) to quantify the deleteriousness of mutations predicted by different methods. Despite the potential for low MAF in neutral/beneficial alleles, we believe this approach provides useful signals for assessing deleterious mutations ^38^. We benchmarked PlantCaduceus against two evolutionary-informed methods, phyloP and phastCons ^29^, as well as GPN ^21^. Both phyloP and phastCons assess evolutionary constraint using multiple sequence alignments and phylogenetic models (**METHODS**), assigning higher scores to conserved regions. We analyzed 4.6 million SNPs in the sorghum TERRA population ^42^ and 9.4 million SNPs from maize Hapmap 3.2.1 population ^46^ and observed that most of the SNPs had neutral zero-shot score, while there was still a heavy tail with negative zero-shot scores (**Fig. 4C**; **Supplemental Fig. 5B**). By defining the top 0.1% as the most deleterious mutations, we observed a significant enrichment in coding regions, as reflected by the high odds ratios in both sorghum (40.70) and maize (42.42) with p-values less than 2.2e-16 (**Supplemental Fig. 6**). We then categorized SNPs into four percentiles based on zero-shot scores: the top 50%, 10%, 1%, and 0.1% most deleterious mutations and observed that all models showed a decreasing average MAF of SNPs in higher percentiles for missense, nonsynonymous, and noncoding SNPs in both sorghum (**Supplemental Fig. 7**) and maize (**Fig. 4D**). Notably, the putative deleterious mutations identified by PlantCaduceus exhibited the lowest average MAF across all percentiles, outperforming GPN and significantly surpassing phyloP and phastCons (**Supplemental Fig. 7; Fig. 4D**). Given the success of protein LMs in predicting deleterious missense mutations ^13,16^, we also incorporated ESM ^12^ as a benchmark. For missense mutations, we found that PlantCaduceus matches the performance of the state-of-the-art protein LM ESM ^12^. At the top 50%, 10%, and 1% percentiles, PlantCaduceus even slightly outperforms ESM in sorghum (**Supplemental Fig. 7**).

However, since GPN is only pre-trained with genomes from eight Brassicales species and specifically designed for mutation effect prediction in Arabidopsis, we further validated PlantCaduceus by analyzing over 10 million mutations from the Arabidopsis 1001 Genomes Project ^47^. Being pre-trained with a broader range of evolutionarily distant genomes, PlantCaduceus effectively captured deleterious mutations in Arabidopsis and slightly outperformed GPN (**Supplemental Fig. 8**). For missense mutations, PlantCaduceus consistently matched the performance of the state-of-the-art protein language model ESM and was nearly competitive with GPN for noncoding mutations.

We further verified if PlantCaduceus could pinpoint known causal deleterious mutations. We collected 19 candidate phenotype-impacting and potentially deleterious mutations identified in homozygous EMS mutants in Arabidopsis ^48^. Among these, 15 mutations were ranked in the top 1% or top 10% by the zero-shot score (**Table 2**), highlighting the zero-shot score of PlantCaduceus can be used for pinpointing causal deleterious mutations. Additionally, PlantCaduceus successfully identified a well-studied causal sweet corn mutation, which derives its characteristic sweetness from the W578R mutation at the *sugary1* (*Su1*) locus ^49^. This mutation disrupts starch metabolism, leading to the accumulation of phytoglycogen, which lowers seedling vigor and reduces germination, ultimately decreasing fitness ^50^. Although GWAS revealed numerous significantly sweet-trait-associated variants on chromosome 4, identifying the exact causal mutations is challenging due to high LD in this low recombination region (**Fig. 5A**). By integrating zero-shot scores from PlantCaduceus with GWAS data (**Fig. 5B-5C**), we successfully identified the W578R mutation as the sole causal variant in this QTL region (**Fig. 5D**). Taken together, these results demonstrate that the PlantCaduceus model effectively pinpoints known causal deleterious mutations, highlighting its potential as a powerful tool for identifying causal variants underlying important agronomic traits.

**Fig 5.**
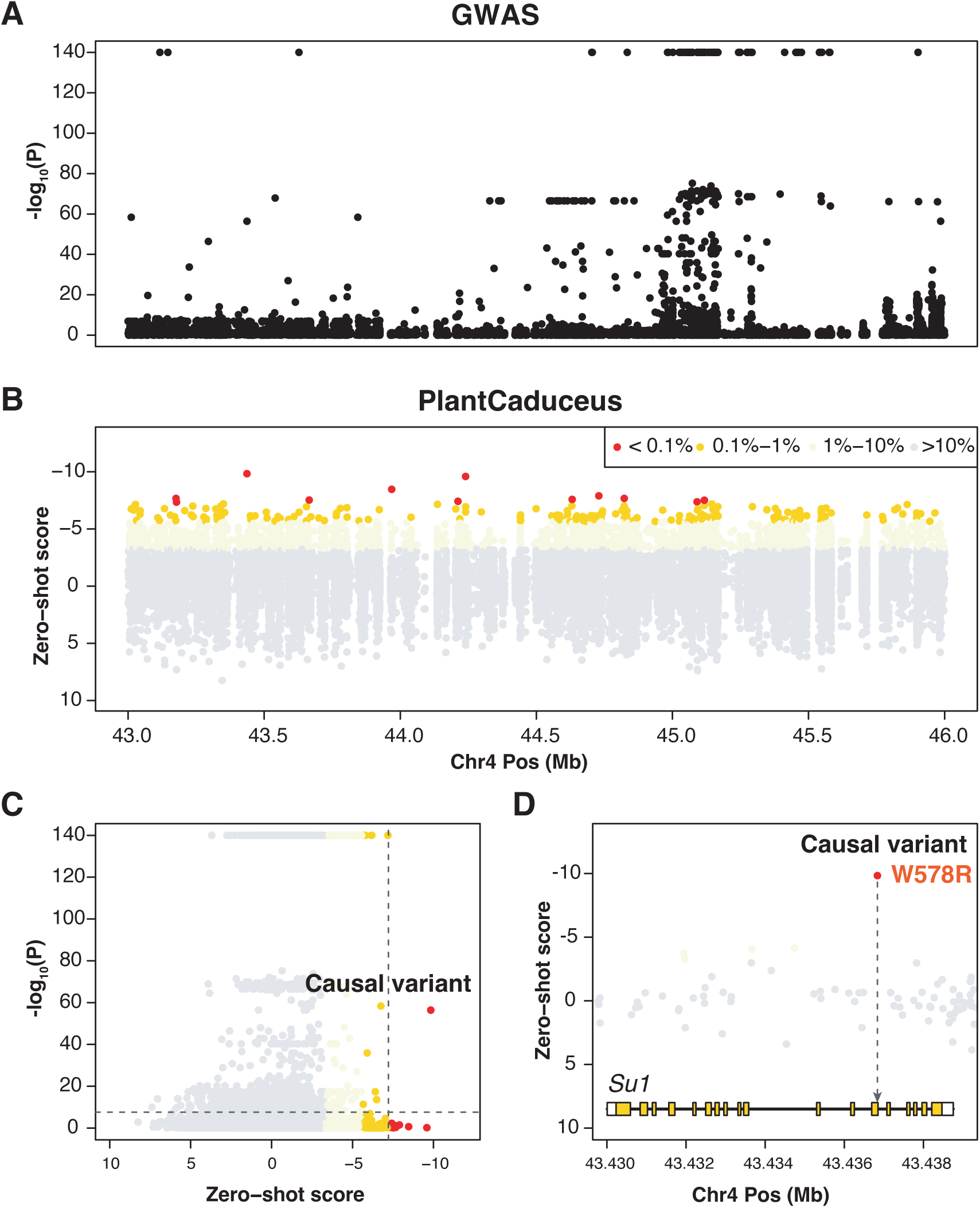
The causal mutation in *Su1* locus. **(A)** Manhattan plot of the sweet corn trait in the region from 43.0 to 46.0 Mb on chromosome 4. **(B)** The zero-shot scores of SNPs in 43.0 to 46.0 Mb in chromosome 4, corresponding to the same region as in **(A)**. **(C)** Scatter plot of zero-shot scores from PlantCaduceus versus −log_10_(P) values from GWAS result. The horizontal dashed line indicates the GWAS significance threshold (Bonferroni’s threshold: 0.05/N; N=2,072,522), and the vertical dashed line marks the top 0.1% percentile of zero-shot scores. (D) Zoomed-in view of the causal variant region and the *Su1* gene structure.

**Table 2.** The zero-shot score of deleterious mutations identified in homozygous EMS mutants.

| Chr | Position | Change | Mutation effect | Phenotype | Zero-shot score | Percentile |
| --- | --- | --- | --- | --- | --- | --- |
| 2 | 14297325 | G>A | Splice change | Univalent chromosomes | -10.344 | Top 1% |
| 3 | 23443192 | C>T | Splice change | Univalent chromosomes | -10.219 | Top 1% |
| 3 | 10277172 | C>T | Splice change | Univalent chromosomes | -9.820 | Top 1% |
| 4 | 5820399 | C>T | Splice change | Univalent chromosomes | -9.547 | Top 1% |
| 3 | 17827101 | G>A | Splice change | Fragmentation | -9.531 | Top 1% |
| 3 | 3248339 | C>T | Premature stop | Univalent chromosomes | -9.125 | Top 1% |
| 3 | 17823207 | G>A | Splice change | Fragmentation | -9.000 | Top 1% |
| 1 | 1298121 | C>T | Splice change | Univalent chromosomes | -8.859 | Top 1% |
| 3 | 17812658 | G>A | Splice change | Fragmentation | -8.719 | Top 1% |
| 5 | 1625685 | G>A | Splice change | Fragmentation | -8.203 | Top 10% |
| 3 | 3246364 | G>A | Splice change | Univalent chromosomes | -7.660 | Top 10% |
| 3 | 3246274 | G>A | Splice change | Univalent chromosomes | -7.406 | Top 10% |
| 5 | 23446256 | G>A | Premature stop | All univalent chromosomes | -6.203 | Top 10% |
| 4 | 16868745 | C>T | Premature stop | Univalent chromosomes | -6.008 | Top 10% |
| 3 | 17824467 | G>A | Missense | Fragmentation | -5.570 | Top 10% |
| 4 | 16868001 | C>T | Premature stop | Univalent chromosomes | -5.141 | Top 50% |
| 3 | 17807938 | G>A | Premature stop | Fragmentation | -4.688 | Top 50% |
| 5 | 26302687 | G>A | Premature stop | Univalent chromosomes | -3.805 | Top 50% |
| 1 | 19964116 | G>A | Start gained | Univalent chromosomes | 0.243 | Top 50% |

## Discussion

Functional annotation of plant genomes is crucial for plant genomics and crop breeding but remains limited by the lack of functional genomic data and accurate predictive models. Here, we introduced PlantCaduceus, a multi-species plant DNA LM pretrained on a curated set of 16 evolutionarily distant Angiosperm genomes, enabling cross-species prediction of functional annotations with limited data. PlantCaduceus leverages Mamba ^28^ and Caduceus ^27^ architectures to support bi-directional, reverse complement equivariant sequence modeling. We demonstrated the superior cross-species performance of PlantCaduceus on five tasks involving transcription, translation, and evolutionary constraint modeling. These results highlight the potential of PlantCaduceus to serve as a foundational model for comprehensively understanding plant genomes.

PlantCaduceus has the potential to accurately annotate any newly sequenced Angiosperm genomes. Unlike supervised deep learning models that easily overfit on limited labeled data, PlantCaduceus demonstrates robust cross-species performance in modeling transcription, translation, and evolutionary constraints, even for species not included in pre-training (**Fig. 2; Supplemental Fig. 2**). This indicates that through self-supervised pre-training on large-scale genomic datasets, PlantCaduceus has captured broad evolutionary conservation and DNA sequence grammar. The cross-species prediction ability of PlantCaduceus can significantly accelerate plant genomics research, aiding initiatives such as the 1000 Plant Genomes Project ^1^ by providing accurate annotations and insights across diverse plant species.

PlantCaduceus offers a more effective approach to estimate deleterious mutations without relying on multiple sequence alignments (MSAs). Deleterious mutations are considered as the genetic basis of heterosis, where hybrids yield more due to the suppression of deleterious recessives from one parent by dominant alleles from the other ^51^. Historically, deleterious mutations have been estimated by generating MSAs ^38,52,53^ and using evolutionary methods such as phyloP and phastCons ^29^. However, the prevalence of transposable elements and polyploidy in plant genomes complicates the genome-wide MSA generation ^54,55^. PlantCaduceus overcomes these challenges by using a masked language modeling strategy to learn conservation from large scale genomic datasets of diverse species. Promisingly, the deleterious mutations prioritized by PlantCaduceus with the zero-shot strategy showed three-fold rare allele enrichment compared to phyloP and phastCons, and our approach is competitive with state-of-the-art protein LM for missense mutations. Furthermore, PlantCaduceus enables pinpointing causal variants from significant GWAS signals, which are usually confounded by LD. These results suggest that PlantCaduceus can be utilized as a powerful tool in crop breeding, enhancing genome-wide deleterious mutation identification, optimizing parental line selection, and promoting hybrid vigor ^51^.

In future work, we plan to incorporate additional plant genomes from diverse lineages, such as gymnosperms, to capture broader evolutionary conservation. Additionally, we plan to pre-train PlantCaduceus with longer context windows, enabling it to capture long-range DNA interactions and better handle tasks benefiting from long-range cis-effects, such as allele-specific expression, chromatin state prediction, and chromatin interaction mapping. Furthermore, it would also be interesting to explore how to better tokenize repetitive sequences in plant genomes. We envision that these approaches will allow us to push the boundaries of what PlantCaduceus can achieve, establishing it as an even more powerful and versatile foundation model for advancing genomic research and facilitating crop improvement.

## Methods

### Pre-training dataset

The pre-training dataset comprises 16 genomes from two distinct clades: eight genomes from the family Poaceae and eight genomes from the order Brassicales (**Supplemental Table 1**). To visualize their relatedness, we subset these taxa from a large phylogeny of seed plants ^56^. The Poaceae species displayed substantial variation in genome size and repetitive sequence content, with the hexaploid wheat genome exhibiting a size of 15 Gbp. For each Poaceae genome, except for Tripsacum, we obtained the genome and corresponding genome annotation and repeat-masked annotation from the Joint Genome Institute (JGI). For the Tripsacum genome, the genome FASTA and annotation files were downloaded from MaizeGDB (https://maizegdb.org/genome/assembly/Td-FL_9056069_6-DRAFT-PanAnd-1.0), and the EDTA tool ^57^ was used to identify repetitive sequences within the genome. Based on the repeat-masked annotation, each genome was softmasked with bedtools ^58^ and subsequently divided into genomic windows of 512 bp with a step size of 256 bp. Each window was assigned to a unique class based on the genome annotation, and all coding sequence regions were selected for pre-training. The remaining genomic regions were then down-sampled to ensure an equal number of CDS regions and noncoding regions. It is important to note that for the hexaploid wheat genome, only subgenome A was utilized to avoid species bias. The Brassicales genomes datasets were acquired from a Hugging Face repository (https://huggingface.co/datasets/songlab/genomes-brassicales-balanced-v1). The validation and testing datasets were randomly selected and constituted 5% of the total dataset.

### Caduceus model architecture and pre-training

We use the recently proposed Caduceus architecture ^27^, which is tailored to DNA sequence modeling. Caduceus is based on the Mamba architecture ^28^, a model which scales to long sequences more efficiently than attention-based methods while maintaining accuracy. Mamba stems from the class of structured state space models (SSMs) ^59^, which are defined by a pair of linear differential equations:

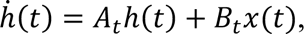

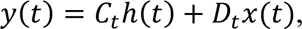

where *x*, *y* ∈ ℝ represent the input and output, respectively, ℎ ∈ ℝ^*n*^ is the state’s hidden representation, and the (potentially time dependent) parameters *A* ∈ ℝ^*n×n*^, *B* ∈ ℝ^*n*×1^, *C* ∈ ℝ^1×*n*^, *D* ∈ ℝ govern the system dynamics. For multi-dimensional inputs and outputs *x*, *y* ∈ ℝ^*R*^, a separate linear system is applied to each of the *d* channels. In practice, using some discretization scheme that is a function of a discrete time parameter Δ, the system is discretized in time, yielding the following:

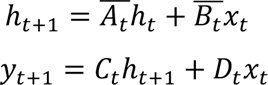

Much of the SSM literature relies on parameters that are fixed in time, allowing for efficient computation during training by means of the convolutional perspective of linear *time invariant* systems ^60^. In contrast to previous SSMs, Mamba enables more expressive models that are *time dependent*, by making the parameters functions of the inputs. This time dependence is crucial in allowing Mamba to overcome the limitations of previous SSMs and rival Transformers ^30^ on sequence modeling tasks across domains. For efficient computation, Mamba employs a parallel algorithm to compute the recurrence relation defined above and an IO-aware implementation that limits potentially bottlenecking memory transfer operations incurred on modern GPU hardware.

To account for upstream and downstream gene interactions, Caduceus employs weight sharing to enable memory-efficient bi-directionality. Finally, Caduceus is designed to consider the reverse complement (RC) symmetry of DNA sequences. This is accomplished by encoding RC equivariance as an inductive bias: the Caduceus language model commutes with the RC operation. Combining these three design decisions, Caduceus has shown promising results when applied to human genome modeling ^27^.

The implementation of RC equivariance in Caduceus entails doubling the number of channels for intermediate representations. At a high level, half the channels are used to encode information about a sequence and the other half are used to encode information about its RC. For downstream tasks in which we fine-tuned a classifier on top of learned embeddings, the labels were invariant to the RC operation, since both DNA strands carry the same label. To account for this, we therefore split embeddings of the Caduceus model along the channel dimension and averaged. This ensures that both a sequence and its RC will have the same final embedding, i.e., we render the embeddings invariant to the RC operation as well.

For the pre-training of PlantCaduceus, each model was trained for 240,000 steps using a Decoupled AdamW optimizer ^61^ with the global batch size of 2,048. The learning rate is 2E-4 with a cosine decay scheduler, and 6% of the training duration was dedicated to warm up. The learning rate decayed to 4E-6 by the end of training. The default BERT ^62^ masking recipe was used with a masking probability of 0.15. For each masked token: (i) there is an 80% probability it will be replaced by a special token ([MASK]), (ii) a 10% probability it will be replaced by a random token, and (iii) a 10% probability it will remain unchanged. Unless otherwise specified, all models were trained using a sequence length of 512 base pairs. A weight decay of 1E-5 was applied throughout the training process.

### TIS, TTS, splice donor and acceptor training, validation and testing dataset generation

To generate high-quality training datasets for translation initiation sites (TIS), translation termination sites (TTS), splice donor sites, and splice acceptor sites, we used the well-annotated model plant genome of Arabidopsis with Araport 11 annotation ^63^. To accurately reflect the inherent imbalance in junction sites prediction, all annotated junction sites were considered as positive observations, while a randomly selected subset of sites (5%) that matched specific appropriate motifs (e.g., ATG for TIS, UAA, UAG, and UGA for TTS, GT for donor splice sites, and AG for acceptor splice sites) were used as negative observations. For each task, the pre-trained model weights were frozen, and XGBoost models (n_estimators=1000, max_depth=6, learning_rate=0.1) were trained using embeddings extracted from the last hidden state of the pre-trained model. To ensure robust model training and validation, chromosome 5 was used for hold-out testing, and the rest of the Arabidopsis genome was used for training.

Given the relatively poor annotation in other species compared to Arabidopsis, we used the BUSCO tool ^64^ to identify 3,236 orthologous genes specific to monocotyledons in *O. sativa*, *S. bicolor* and *Z. mays* and 2,326 orthologous genes specific to eudicotyledons in *G. hirsutum* and *G. max* to generate reliable testing datasets in other species. This approach ensures that the selected annotated genes are highly conserved and likely to be correctly annotated, mitigating the issue of inaccurate performance evaluations. Specifically, BUSCO was utilized to scan the annotated protein isoforms, and only complete BUSCO genes were considered as true positives. For those BUSCO genes with multiple transcripts, we selected the longest transcript to avoid sequence redundancy in the testing dataset. Subsequently, BUSCO gene/transcript-supported junction sites were used as positive examples for their respective tasks. To generate negative sites, all sites within the BUSCO genes that matched appropriate motifs (e.g., ATG for TIS, TAA, TAG, and TGA for TTS, GT for donor splice sites, and AG for acceptor splice sites) but were not part of any annotated gene models were used as true sites. Sites belonging to alternate transcripts were excluded to avoid ambiguity. Furthermore, to expand the negative observations and capture a broader range of non-junction sites, we included sites in the intergenic regions flanking the BUSCO genes that matched the appropriate junction motifs. By incorporating both genic and intergenic sites from the BUSCO gene set as negatives, we created an extremely imbalanced testing dataset to reflect the real-world scenario of junction site prediction (**Supplemental Table 3**).

### Evolutionary constraint estimation

The evolutionary constraint was estimated primarily within the Andropogoneae tribe, a large clade of grasses comprising approximately 1,200 species that descended from a common ancestor approximately 18 million years ago ^39^. In this analysis, 34 genomes from Andropogoneae and the rice genome were used to estimate the evolutionary constraint. Due to the substantial transposable element (TE) content in these genomes, AnchorWave, a sensitive genome-to-genome alignment tool ^54^, was used to align the 35 genomes to the sorghum reference genome using the parameters “-R 1 -Q 1”. Following the alignments to the sorghum reference genome, we counted the number of identities, SNPs, and coverages (**Supplemental Fig. 3**). Then the fine-tuned labels were generated based on per-site identity and coverage (**Fig. 3A**). Conserved sites were defined as having an identity greater than 34, while neutral sites were defined as having an identity of 15 or less and coverage of at least 34. Sites with low coverage were excluded due to their potential ambiguity. Given the large size of the training dataset, only 5% of conserved sites were randomly selected for training, and an equivalent proportion of neutral sites was also randomly selected. Sites from chromosomes 1 to 9 were used for training, while those from chromosome 10 were used for validation. To generate the testing dataset in maize, the maize reference genome B73 was used. Then, using the same approach, genome-wide evolutionary constraints were generated by aligning 35 genomes to the maize reference genome with AnchorWave, using the parameters “-R 1 -Q 2,” except for Tripsacum clades. For Tripsacum and maize, which share the most recent whole genome duplication, we used “-R 1 -Q 1”.

### phyloP and phastCons calculation

With the same 34 genomes from Andropogoneae, we generated pairwise genome-to-genome alignments using Cactus ^65^, a multiple genome alignment tool that uses a progressive alignment strategy. The neutral model was calculated from fourfold degenerate coding sites across the entire genome. The resulting alignments were then analyzed using PHAST ^29^ to quantify evolutionary conservation with phyloP conservation scores – using the SPH scoring method (--method SPH) and CONACC mode (--mode CONACC) – and phastCons scores.

### In silico mutagenesis

All potential mutations in the genic regions and 1 kb flanking regions of maize and sorghum chromosome 8 were generated and annotated using the Ensembl Variant Effect Predictor (VEP) local API ^44^, with the upstream/downstream parameter set to 1,000 to classify variants as either upstream or downstream. For intergenic variants, we randomly sampled 100,000 SNPs from the intergenic regions across chromosome 8 to ensure more even coverage of the entire chromosome. For each variant type, we randomly sampled 100,000 mutations and calculated zero-shot scores.

### Genome-wide association study for sweet phenotype

To perform a GWAS for the sweet phenotype, we used a subset of genotypes from the Hapmap 3.2.1 population ^46^, where sweet phenotype data is available ^66^. This subset consists of 272 diverse inbred lines with recorded sweet phenotype data. We coded starchy corn as 0 and sweet corn as 1, with 266 entries in the first category and 6 in the second. To map the sweet phenotype, we utilized a model specifically designed to account for population structure: y = Xβ1 + 5 global PCs + e. The methods for GWAS followed those outlined in Kpaipho-Burch et al ^67^. Briefly, the five global principal components (PCs) were derived from 66,527 SNPs across 3,545 diverse inbred lines, and the SNPs from 272 inbred lines were then rotated to such PCs. The selected SNPs had no missing data across three maize populations, ensuring effective control for population structure and kinship. This approach also reduced computational time compared to mixed linear models while maintaining consistent trait mapping across populations.

### GPN, custom GPN, AgroNT and NT-v2 baselines

To comprehensively evaluate our foundation model’s performance, four foundation models including GPN ^21^ (https://huggingface.co/songlab/gpn-brassicales), custom GPN, AgroNT ^23^ (https://huggingface.co/InstaDeepAI/agro-nucleotide-transformer-1b) and NT-v2 ^24^ (https://huggingface.co/InstaDeepAI/nucleotide-transformer-v2-500m-multi-species) were used as baselines for various tasks. GPN is a convolutional DNA LM pre-trained on eight genomes of Arabidopsis and seven other species from the Brassicales order. However, since GPN was pre-trained with only eight evolutionarily close species and has only 65M parameters and most of the tasks in this paper focus on evaluation in crops, we re-trained a custom GPN with 130M parameters using 50 convolutional layers and the same dataset as PlantCaduceus for a fair comparison. The other hyperparameters were kept identical to the original GPN (**Supplemental Table 6**). In contrast, AgroNT ^23^ is a transformer-based ^30^ language model with 1 billion parameters, pre-trained on 48 plant genomes. NT-v2 ^24^, is a non-plant multi-species transformer model pre-trained on 850 genomes excluding plant species. These models employ different tokenization strategies: GPN uses single-nucleotide tokenization, while AgroNT and NT-v2 use 6-mer tokenization. To ensure a fair comparison, we extracted the middle token embeddings for GPN and the middle k-mer token embeddings for AgroNT and NT-v2.

### Supervised CNN+LSTM baseline

To establish a fair comparison between our DNA LM and existing supervised models, which are primarily trained on human data, we used the DanQ model architecture ^35^ as the supervised baseline. DanQ is a hybrid convolutional and recurrent neural network specifically designed for predicting the function of DNA sequences. It has demonstrated impressive performance in predicting chromatin states in plant species, making it a suitable choice for our comparative analysis ^68^. For each task, the CNN+LSTM model was trained from scratch using one-hot encoded DNA sequences as input. The Adam optimizer with a learning rate of 0.01 was employed for model optimization. The batch size was set to 2,048. Early stopping with a patience of 20 steps was implemented.

## Data availability

The pre-training genomes are available at: https://huggingface.co/datasets/kuleshov-group/Angiosperm_16_genomes. All datasets used for fine-tuning are available at Hugging Face: https://huggingface.co/datasets/kuleshov-group/cross-species-single-nucleotide-annotation

## Code availability

The pre-trained models, along with documentation on how to use them, are available at Hugging Face: https://huggingface.co/collections/kuleshov-group/plantcaduceus-512bp-len-665a229ee098db706a55e44a. The pre-training and fine-tuning codes are available at GitHub: https://github.com/kuleshov-group/PlantCaduceus

## Acknowledgments

This work is funded by the USDA-ARS, NSF PanAnd grant (#1822330), NSF CAREER grant (#2145577) and NIH MIRA grant (#1R35GM151243-01). We thank Edgar Marroquin (Cornell University) for discussing fine-tuning tasks, Travis Wrightsman (Cornell University) for providing DanQ code, Arun S. Seetharam and Matthew B Hufford (Iowa State University) for sharing Andropogoneae assemblies, Merritt Khaipho-Burch (Cornell University) for sharing the liftover version HapMap3 VCF file, Sara Miller (Cornell University) for helpful comments and all members of the E.S.B. laboratory (Cornell University) for helpful discussions. We would also like to thank the SCINet project, the AI Center of Excellence of the USDA Agricultural Research Service (0201-88888-003-000D and 0201-88888-002-000D) and MosaicML for providing compute resources for pre-training and fine-tuning experiments.

## Author contributions

J.Z., A.G., E.S.B., and V.K. designed the research; J.Z., A.B., Z.-Y.L., Z.R.M., A.S., M.C.S. and C.R. curated the data; J.Z., A.G., Y.S., and Z.R.M. pre-trained models; J.Z., A.B., Z.-Y.L., and Z.R.M. performed fine-tuning tasks; J.Z., A.B., Z.-Y.L., W.-Y.L., and Z.R.M. analyzed results; J.Z. wrote the manuscript and all authors edited the manuscript.

