## Supplemental Figures for "Cross-species modeling of plant genomes at single nucleotide resolution using a pre-trained DNA language model"

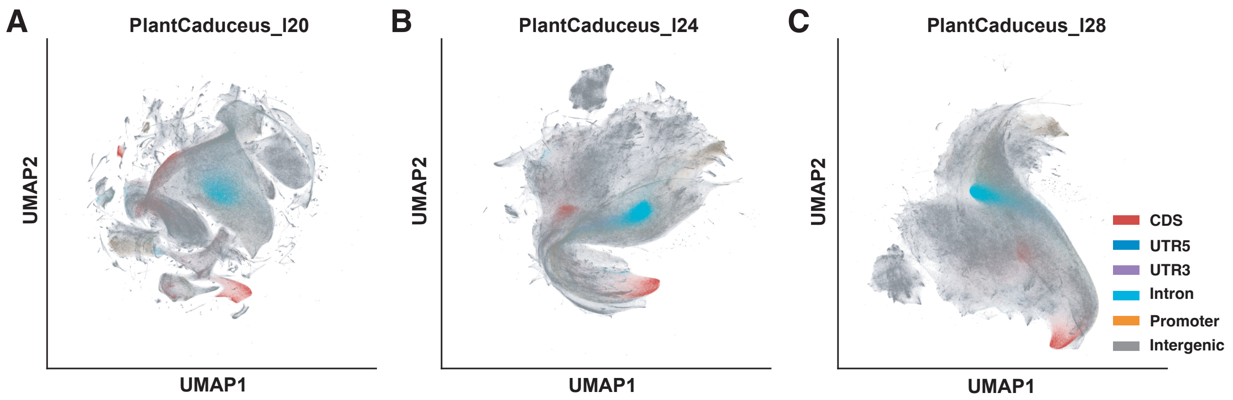


**Supplemental Fig 1. Unsupervised clustering of genomic windows.** UMAP visualization of embeddings from PlantCaduceus_l20 **(A)**, PlantCaduceus_l24 **(B)**, and PlantCaduceus_l28 **(C)** averaged over non-overlapping 100-bp windows along the sorghum genome. PlantCaduceus_l20, PlantCaduceus_l24, and PlantCaduceus_l28 represent PlantCaduceus models with 20 layers, 24 layers, and 28 layers, respectively.

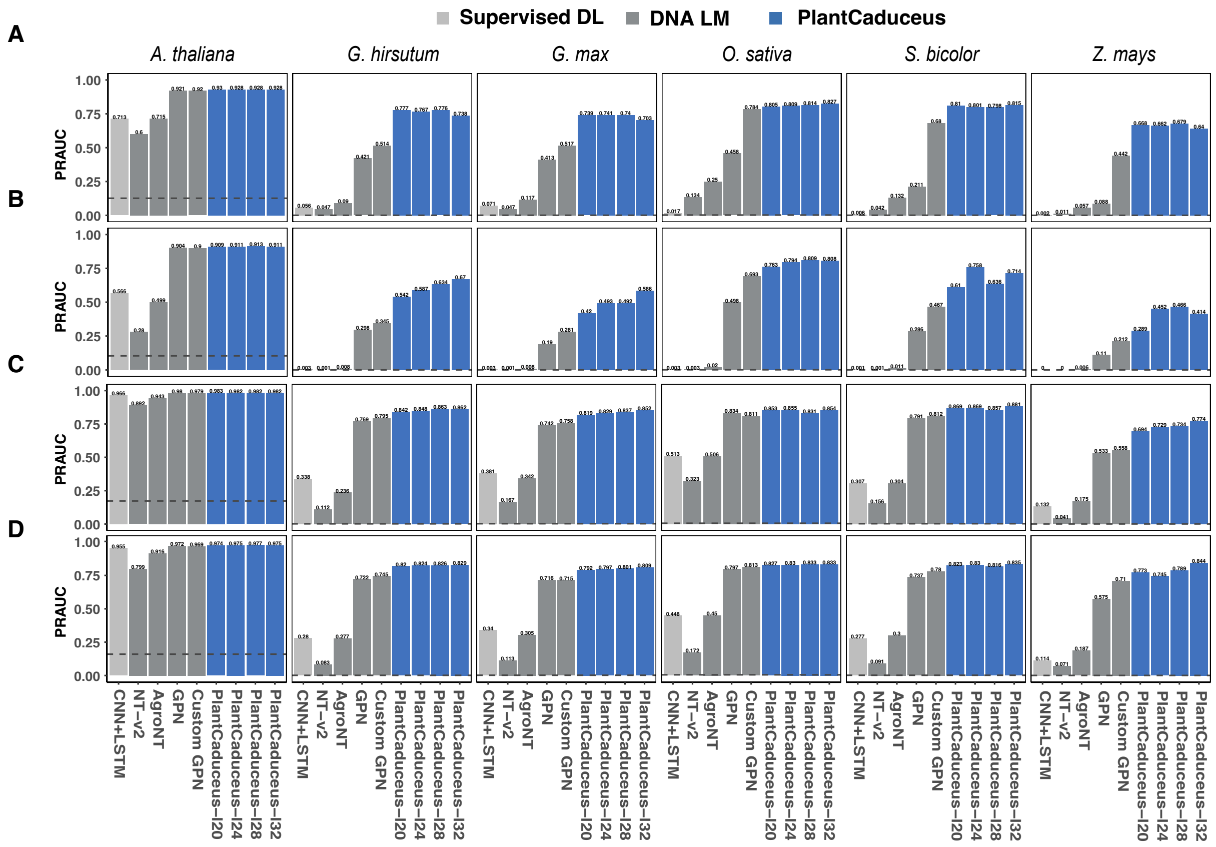


**Supplemental Fig 2. Performance evaluation of TIS, TTS, splice donor and acceptor prediction tasks.** Bar plots displaying the PRAUC scores of different models across various species: *A. thaliana* (with-species), *G. hirsutum*, *G.max*, *O.sativa*, *S.bicolor* and *Z. mays* for four tasks: TIS **(A)**, TTS **(B)**, donor **(C)**, and acceptor **(D)**. The blue bars represent our four PlantCaduceus models with varying layers. The gray bars denote three DNA language models: NT-v2, AgroNT, GPN, and Custom GPN (pre-trained with the same data as PlantCaduceus). The light gray bars represent a traditional supervised model, which is a hybrid of CNN and LSTM. The gray dashed line in each panel represents the baseline for each dataset, which corresponds to the negative sample ratio.

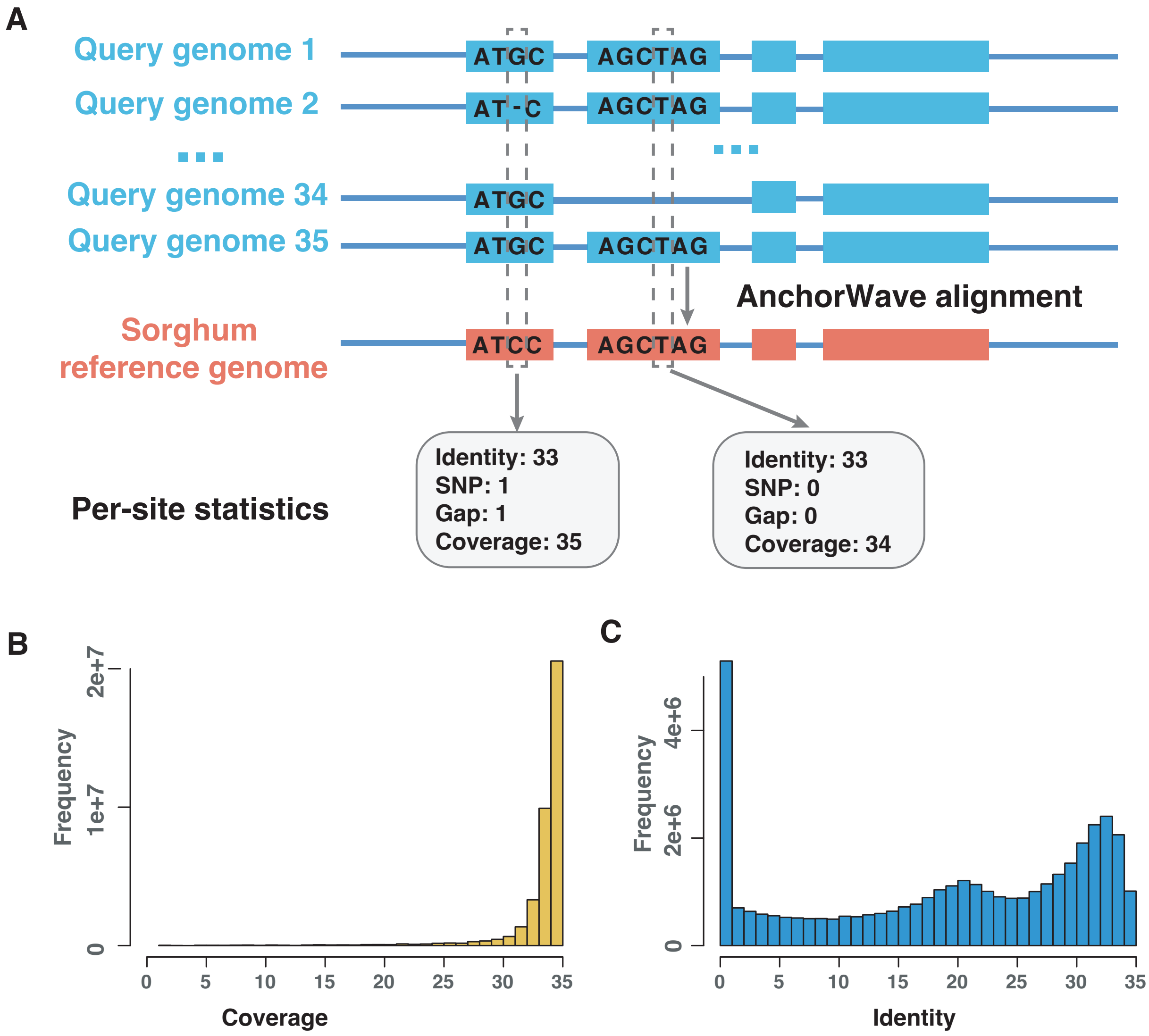


**Supplemental Fig 3. Evolutionary constraint estimation.** **(A)** Illustration of the evolutionary conservation estimation process. **(B)** Distribution of coverage in coding regions. **(C)** Distribution of identity in coding regions.

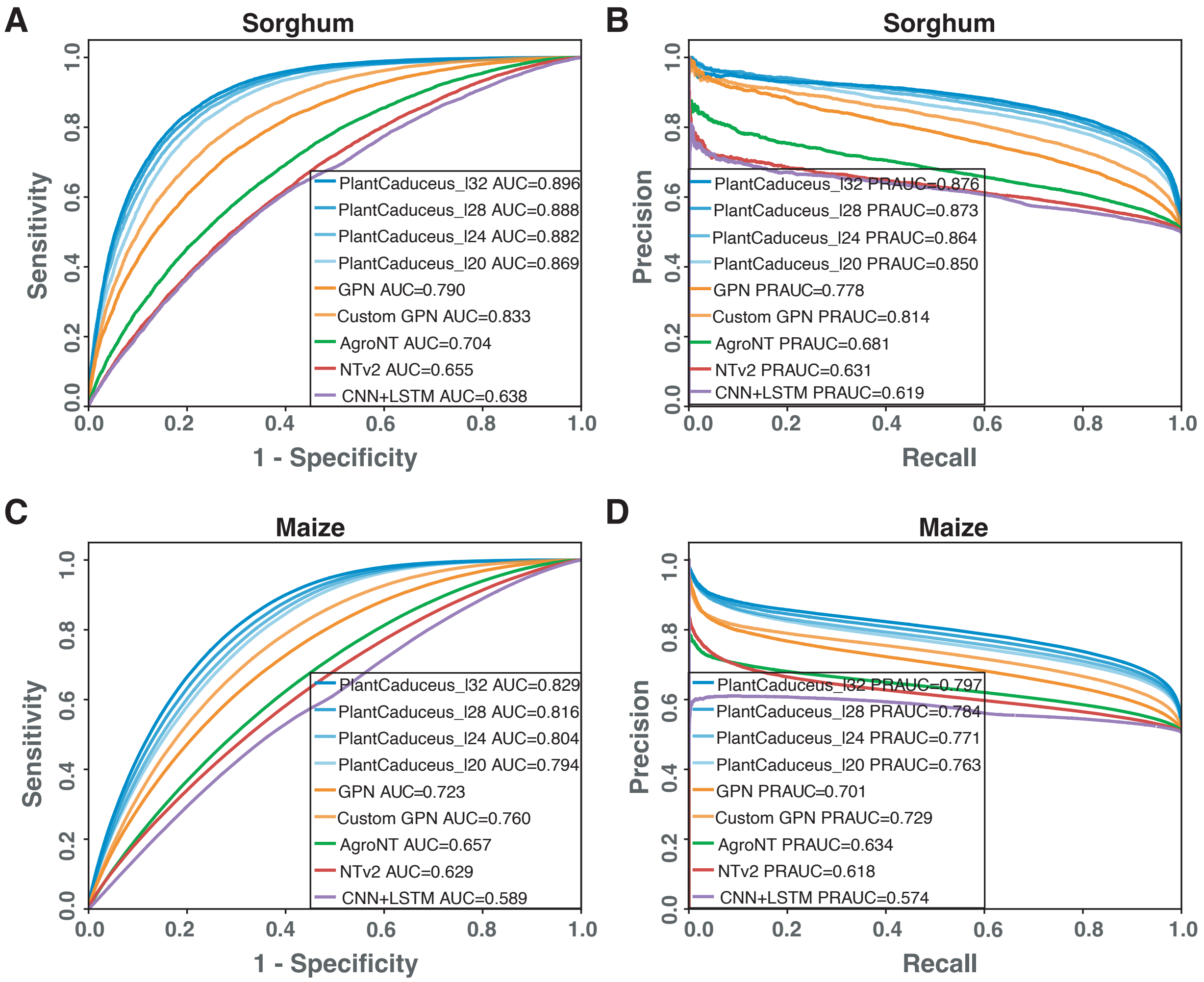


**Supplemental Fig 4. Evolutionary constraint performance evaluation. (A)** Receiver operating characteristic (ROC) and **(B)** precision-recall (PR) curves of different models in sorghum. **(C)** ROC and **(D)** PR curves of transferring different models trained in sorghum to unseen maize data.

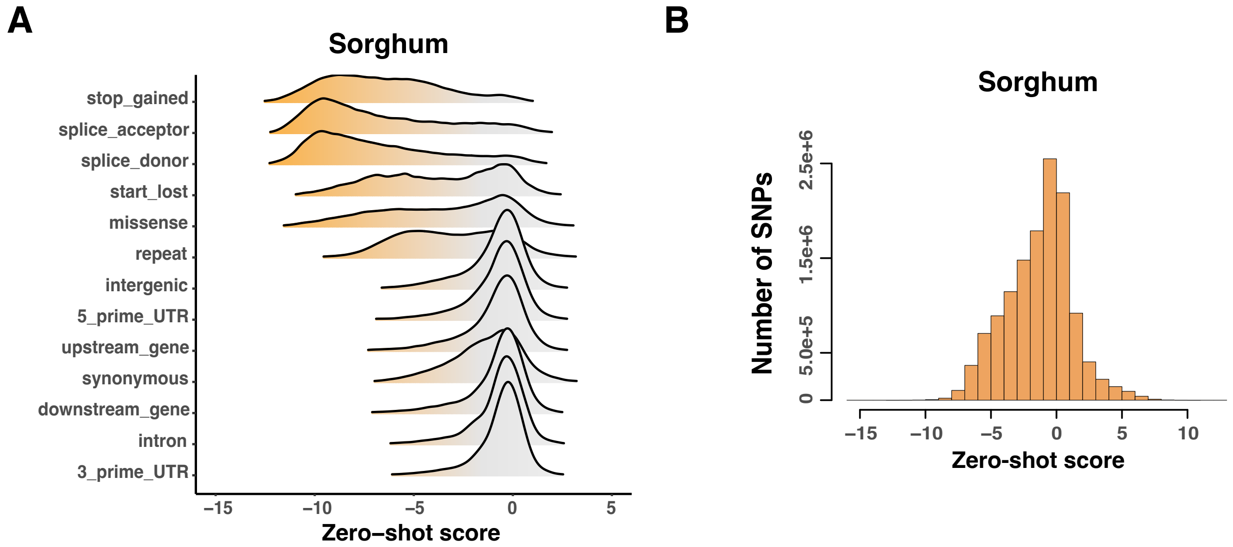


**Supplemental Fig 5. Deleterious mutations identification in Sorghum. (A)** The zero-shot score distribution of different types of variants generated by in silico mutagenesis in sorghum. **(B)** The zero-shot score distribution of 4.6M SNPs in the sorghum TERRA population.

**A** Sorghum

| | Top 0.1% deleterious SNPs | Other SNPs |
|---|---|---|
| **CDS** | 3,700 | 443,259 |
| **Not CDS** | 840 | 4,097,716 |

Odds ratio = 40.70; p-value < 2.2e-16

**B** Maize

| | Top 0.1% deleterious SNPs | Other SNPs |
|---|---|---|
| **CDS** | 7,136 | 658,111 |
| **Not CDS** | 2,222 | 8,706,658 |

Odds ratio = 42.42; p-value < 2.2e-16

**Supplemental Fig 6. CDS enrichment for predicted deleterious mutations in sorghum and maize.** Contingency table and odds ratio showing the enrichment of putative deleterious SNPs in CDS regions for sorghum **(A)** and maize **(B)**.

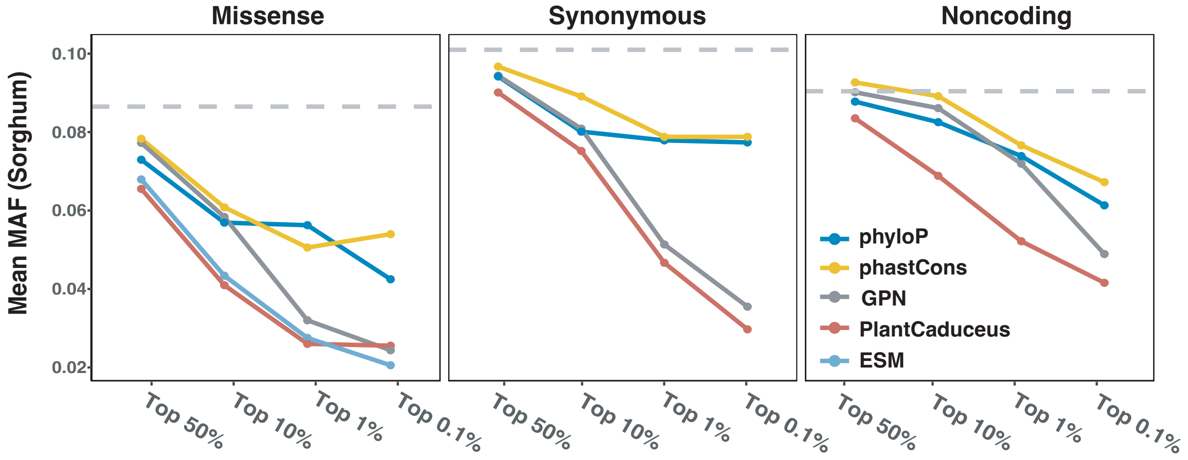


**Supplemental Fig 7.** The MAF of putative deleterious mutations prioritized by different models in sorghum.

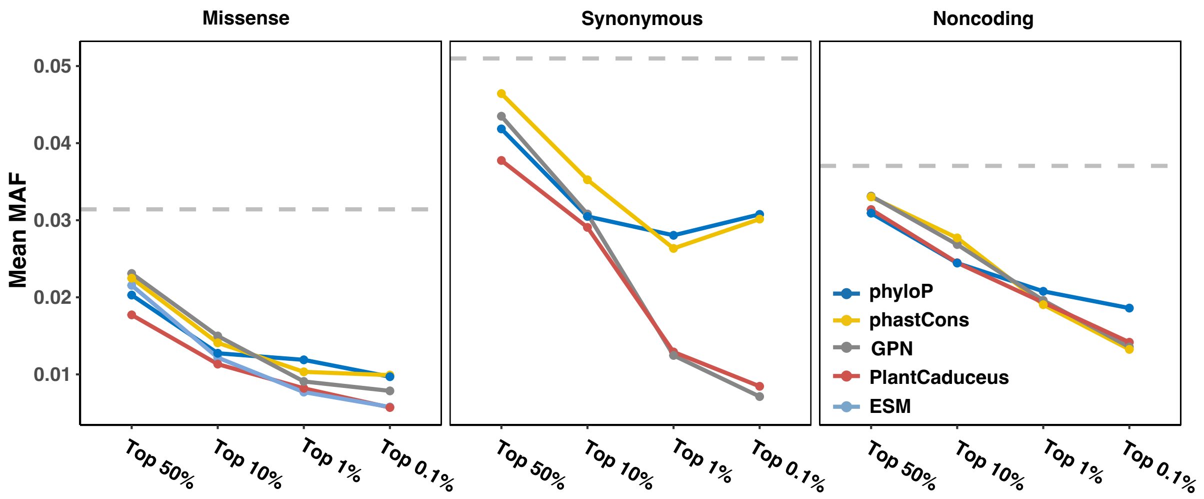


**Supplemental Fig 8.** The MAF of putative deleterious mutations prioritized by different models in Arabidopsis.
